# Impacts of Biostimulation and Bioaugmentation on Woodchip Bioreactor Microbiomes

**DOI:** 10.1101/2022.10.06.511109

**Authors:** Hao Wang, Gary W. Feyereisen, Ping Wang, Carl Rosen, Michael J. Sadowsky, Satoshi Ishii

## Abstract

Woodchip bioreactors (WBRs) are used to remove nutrients, especially nitrate, from subsurface drainage. The nitrogen removal efficiency of WBRs, however, is limited by low temperature and availability of labile carbon. Bioaugmentation and biostimulation are potential approaches to enhance nitrate removal of WBRs under cold conditions, but their effectiveness is still unclear. Here, we clarified the effects of bioaugmentation and biostimulation on the microbiomes and nitrate removal rates of WBRs. As a bioaugmentation treatment, we inoculated WBR-borne cold-adapted denitrifying bacteria *Cellulomonas cellasea* strain WB94 and *Microvirgula aerodenitrificans* strain BE2.4 into the WBRs located at Willmar, MN, USA. As a biostimulation treatment, acetate was added to the WBRs to promote denitrification. Woodchip samples were collected from multiple locations in each WBR before and after the treatments and used for the microbiome analysis. The 16S rRNA gene amplicon sequencing showed that the microbiomes changed by the treatments and season. The high-throughput quantitative PCR for nitrogen cycle genes revealed a higher abundance of denitrification genes at locations closer to the WBR inlet, suggesting that denitrifiers are unevenly present in WBRs. In addition, a positive relationship was identified between the abundance of *M. aerodenitrificans* strain BE2.4 and those of *norB* and *nosZ* in the WBRs. Based on generalized linear modeling, the abundance of *norB* and *nosZ* was shown to be useful in predicting the nitrate removal rate of WBRs. Taken together, these results suggest that the bioaugmentation and biostimulation treatments can influence denitrifier populations, thereby influencing the nitrate removal of WBRs.

**IMPORTANCE:** Nitrate pollution is a serious problem in agricultural areas in the U.S. Midwest and other parts of the world. Woodchip bioreactor is a promising technology that uses microbial denitrification to remove nitrate from agricultural subsurface drainage, although the reactor’s nitrate removal performance is limited under cold conditions. This study showed that the inoculation of cold-adapted denitrifiers (i.e., bioaugmentation) and the addition of labile carbon (i.e., biostimulation) can influence the microbial populations and enhance the reactor’s performance under cold conditions. This finding will help establish a strategy to mitigate nitrate pollution.

## INTRODUCTION

Subsurface drainage systems are commonly used in agricultural areas in the Upper Midwest in the U.S. as well as northern Europe to optimize soil conditions for plant roots, provide benefits to crop yield, and improve field trafficability during the planting and harvesting season (Zucker and Brown, 1998); however, at the same time, they unintentionally transport nutrients, especially nitrate, to receiving natural waterbodies, causing various detrimental effects on the ecosystem such as eutrophication (Huffman et al., 2013; Porter et al., 2015).

Various methods have been used to remove nitrate from agricultural drainage including wetlands, managed drainage systems, and denitrification bioreactors (Dinnes et al., 2002). Among those, end-of-the-pipe denitrification bioreactor is a promising technology to treat subsurface drainage with minimum impacts on farmland (Schipper et al., 2010; Bednarek et al., 2014; Addy et al., 2016; Christianson et al., 2020). Woodchip bioreactor (WBR) is the most commonly used denitrification bioreactor, in which denitrifying organisms use woodchips as a source of carbon and electron donor to reduce nitrate to dinitrogen gas (Fowdar et al., 2015). WBR has been shown to be effective for removing nitrate from agricultural drainage (Schipper et al., 2010; Addy et al., 2016). However, the nitrate removal rate (NRR) of WBR is limited when the water temperature is low most likely due to limited microbial activity (David et al., 2016; Feyereisen et al., 2016; Hoover et al., 2016; Roser et al., 2018).

Potentially useful approaches to improve nitrate removal efficiency of WBR are biostimulation and bioaugmentation. Biostimulation is the enhancement of microbial activity by adding respiration substrate or nutrients or optimizing environmental conditions (e.g., oxygen), whereas bioaugmentation is the inoculation of microorganisms capable of carrying out the desired bioremediation reaction (Tyagi et al., 2011). Previous laboratory experiments showed that the addition of acetate to woodchip column reactors increased nitrate removal rate at cold conditions (5.5°C) (Roser et al., 2018). However, microbial analysis was not conducted, and therefore, it is unknown what kinds of microbes were enriched by the acetate addition. For the purpose of bioaugmentation, we previously identified and isolated denitrifiers that were active at relatively low temperatures (15°C) from WBR (Jang et al., 2019; Anderson et al., 2020). Some of them can breakdown cellulose, a major component of woodchips, and therefore, can provide more labile carbon to the environment (Jang et al., 2019). By inoculating these cold-adapted denitrifiers, it might be possible to enhance nitrate removal at cold conditions (Jéglot et al., 2021, 2022).

We conducted a field-scale biostimulation and bioaugmentation campaign to enhance NRR in a WBR in Minnesota with each treatment replicated two times. In this campaign, biostimulation (i.e., the addition of acetate), bioaugmentation (i.e., the addition of cold-adapted denitrifiers), and an nonamended control were prepared and monitored over 1.5 years. Engineering and water quality aspects of this campaign have been reported by Ghane et al. (2019) and Feyereisen et al. (2022), respectively. The current study focuses on the microbial aspects of this campaign. Specifically, the objectives of this study were to (i) compare the abundance of nitrogen cycle genes before and after the biostimulation/bioaugmentation treatments and throughout the experimental period, (ii) determine the microbiome changes in response to the treatment, and (iii) identify the microbial and environmental factors influencing NRR of WBR. We used a high-throughput nitrogen cycle gene quantification tool called NiCE chip (Oshiki et al., 2018), the 16S rRNA gene amplicon sequencing, and statistical modeling to meet these objectives.

## RESULTS

### Nitrate removal performance of WBR

Nitrate-N removal rates were significantly higher for the biostimulation treatment than the bioaugmentation and control treatments for the Spring 2017, 15.0, 5.8, and 4.4 mg N m^-3^ d^-1^ (*P* = 0.029), and Fall 2017 campaigns, 5.6, 3.9, and 4.1 mg N m^-3^ d^-1^ (*P* = 0.095), respectively (Feyereisen et al., 2022). Nitrate-N load removal was also higher for biostimulation than for bioaugmentation and control treatments for the these two campaigns: 65, 21, and 17%, respectively, for Spring 2017, and 31, 20, and 16%, respectively, for Fall 2017. There were dates after inoculation wherein effluent nitrate-N concentrations for bioaugmentation were reduced relative to the control, but this effect was not sustained. Nitrate-N removal rates for Spring 2018 were not significantly different among treatments (*p*=0.54).

### Nitrogen cycle gene abundance

Among the 36 assays used in the NiCE chip, 24 assays produced successful amplifications on more than half of the samples and had a standard curve with at least three points and an r^2^ value >0.95, and therefore, were used for further analysis. These genes include genes for nitrification (*amoA, hao*, and *nxrB*), denitrification (*napA, nirK, nirS, norB*, and *nosZ* clade I & II), and anaerobic ammonium oxidation (anammox; *hdh*) (Table S3). Assays used to measure *narG* (denitrification), *hzs* (anammox), *nrfA* (DNRA), and *nifH* (nitrogen fixation) genes failed to amplify and therefore were removed from further analysis. Overall, the heatmap generated using the NiCE chip data showed higher relative abundances of denitrification genes than that of nitrification genes in woodchip bioreactor (Fig. S1). The heatmap of denitrification genes showed higher relative abundances in samples collected from port #2, which is near the bed inlet, compared to samples collected from ports #4 and #5, which are closer to the bed outlet (Fig. 1). This was also supported by paired t-test. Of the twelve assays targeting denitrification genes, five of them (napA_v66, nirK_FlaCu, norB_qnorB2F, norB_2, nosZ_1F) showed higher abundance in samples collected from port #2 than in those collected from port #5 (*p* < 0.05). This suggest that denitrifying bacteria were unevenly distributed along the WBR and there may be some active denitrification zone in WBR: i.e., denitrifying bacteria may be more abundant in locations closer to the inlet where nitrate concentration was the highest (Feyereisen et al., 2022).

**Figure 1.**
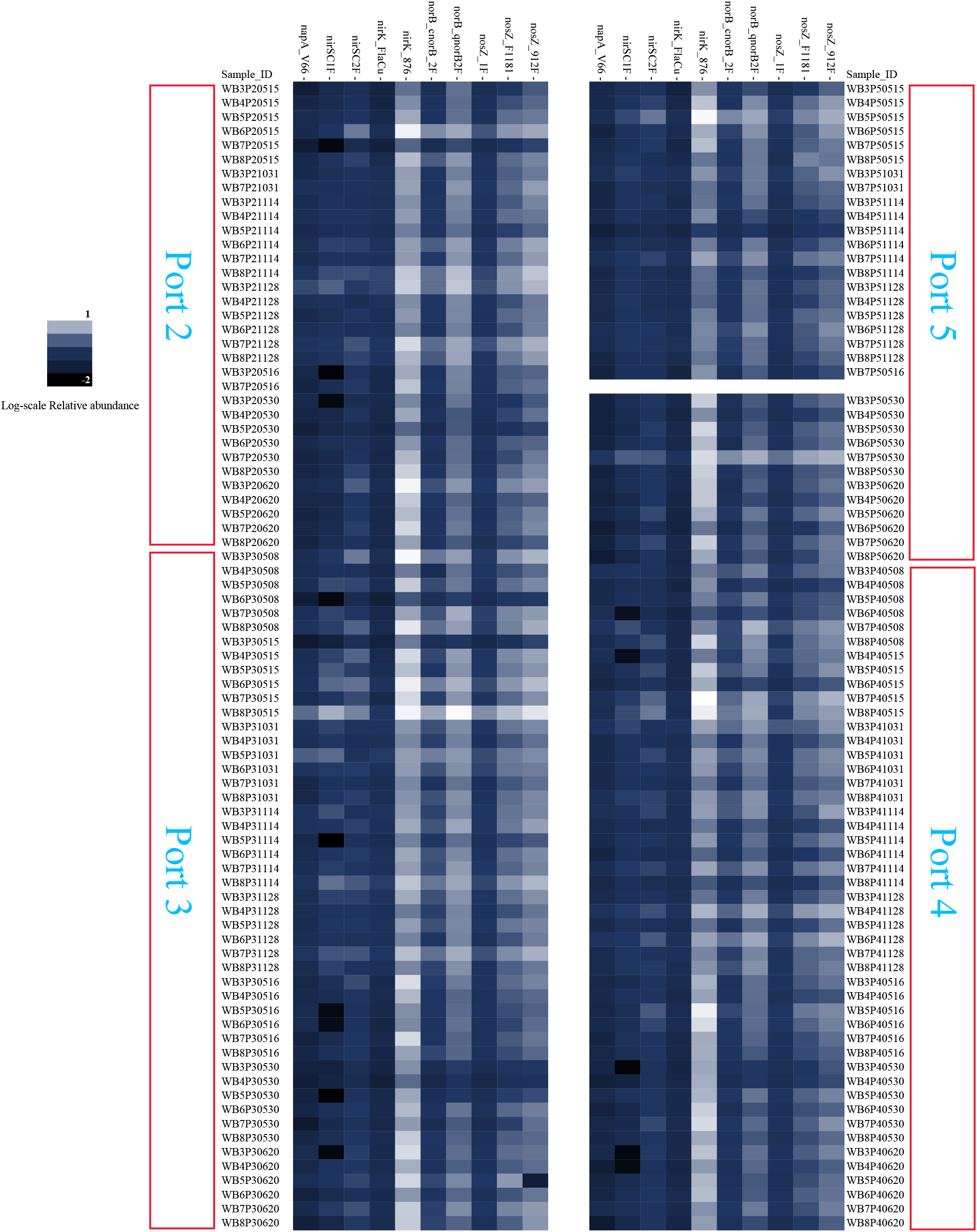
Heatmap showing the log-scale relative abundances of denitrification genes measured by the Nitrogen Cycle Evaluation (NiCE) chip. Samples were grouped by the port number.

The relationship between the environmental variables and the N-cycle associated genes were visualized using a correlation plot (Fig. 2). Strong positive correlations were seen between all of the nitrogen cycle related genes analyzed. The abundance of *Microvirgula aerodenitrificans* strain BE2.4 was also positively correlated with *norB* and *nosZ* abundances. 16S rRNA gene copy number was positively correlated with temperature but almost all of the N cycle-related genes normalized by using the 16S rRNA gene copy number as the denominator were negatively correlated to temperature. This suggests that higher temperatures increased the overall bacterial biomass more so than N cycle related genes. In addition, the port number was negatively related to almost all of the nitrogen cycle related genes, especially the genes involved in the denitrification process, further supporting the uneven distribution of denitrification genes within WBR.

**Figure 2.**
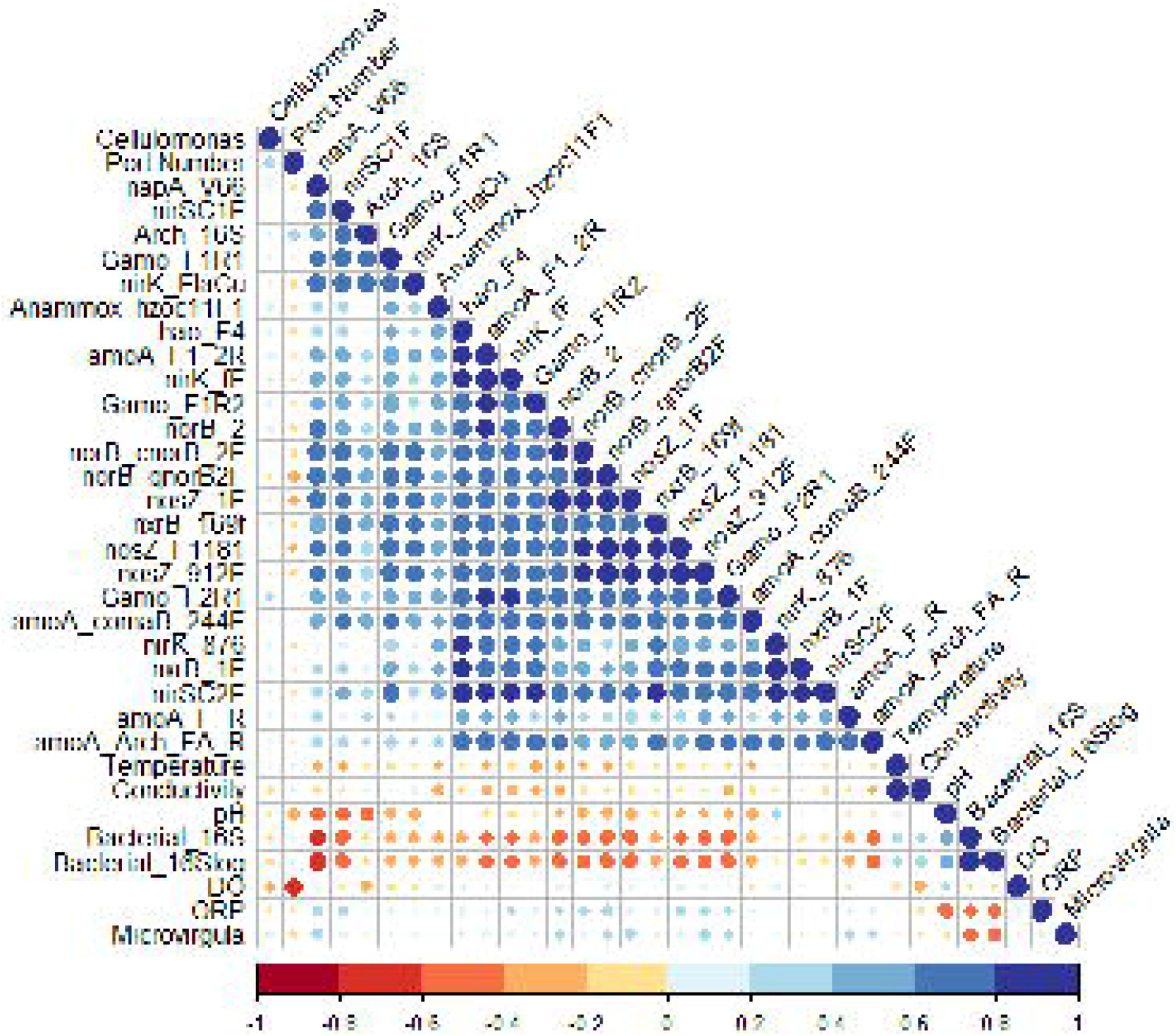
Correlation plots of the NiCE chip measurements, abundance of the inoculated bacteria, and environmental variables. The color intensity and the size of the circle are proportional to Spearman’s ρ value.

Based on these findings, we selected port #2 as the denitrification hot spot in WBR and used data from this port for the GLM analysis. Specifically, the data collected 3 from port #2 on different dates were used to predict the weekly average of NRR (g N m^-3^ d^-1^) for each WBR. The GLM results showed that Assay norB_2 and Assay nosZ_912F have statistically significant (*p* < 0.10) coefficients in explaining the changes in the NRR. Both *norB* and *nosZ* are involved in the denitrification process, and the GLM model showed that they both have a positive relationship with the NRR.

### WBR microbiomes

The 16S rRNA gene amplicon sequencing was used to analyze the microbiome structures in WBR with different treatments. At least 5000 sequence reads per sample were obtained from 158 of the 169 samples. Samples with less than 5000 reads were removed from the downstream analyses. For the 158 samples, the mean and median sequencing depths were 13,153 and 13,275 reads, respectively (Fig. S2).

Alpha-diversity of the WBR microbiome was assessed by calculating Shannon index values. While mean Shannon index value of samples collected from the bioaugmentation bioreactors (bed #3 and #7) was slightly elevated compared to that of samples collected from the control and biostimulation beds (Fig. S3); there was no statistically significant difference among treatment groups (*p*=0.262 by ANOVA). In addition, no statistical difference was seen in samples collected at different time points throughout the sampling period (from May 2017 to June 2018).

The ordination technique (PCoA) was used to visualize the β-diversity of the WBR microbiomes. On the PCoA plot (Fig. 3), samples collected in spring (Spring 2017 and Spring 2018) were clustered together and relative distances plotted from those collected in Fall 2017. PERMANOVA test showed that there was a statistically significant difference (*p* <0.001) between the microbiomes of samples collected in different seasons and years. The PERMANOVA test showed that the temporal change (different sampling dates from May 2017 to June 2018) accounts for 17.1% of the difference between the WR microbiomes, indicating that the season may have a substantial impact on the microbiomes. On the other hand, treatments accounted for only 2.5% of the difference between the WBR microbiomes (*p*=0.002).

**Figure 3.**
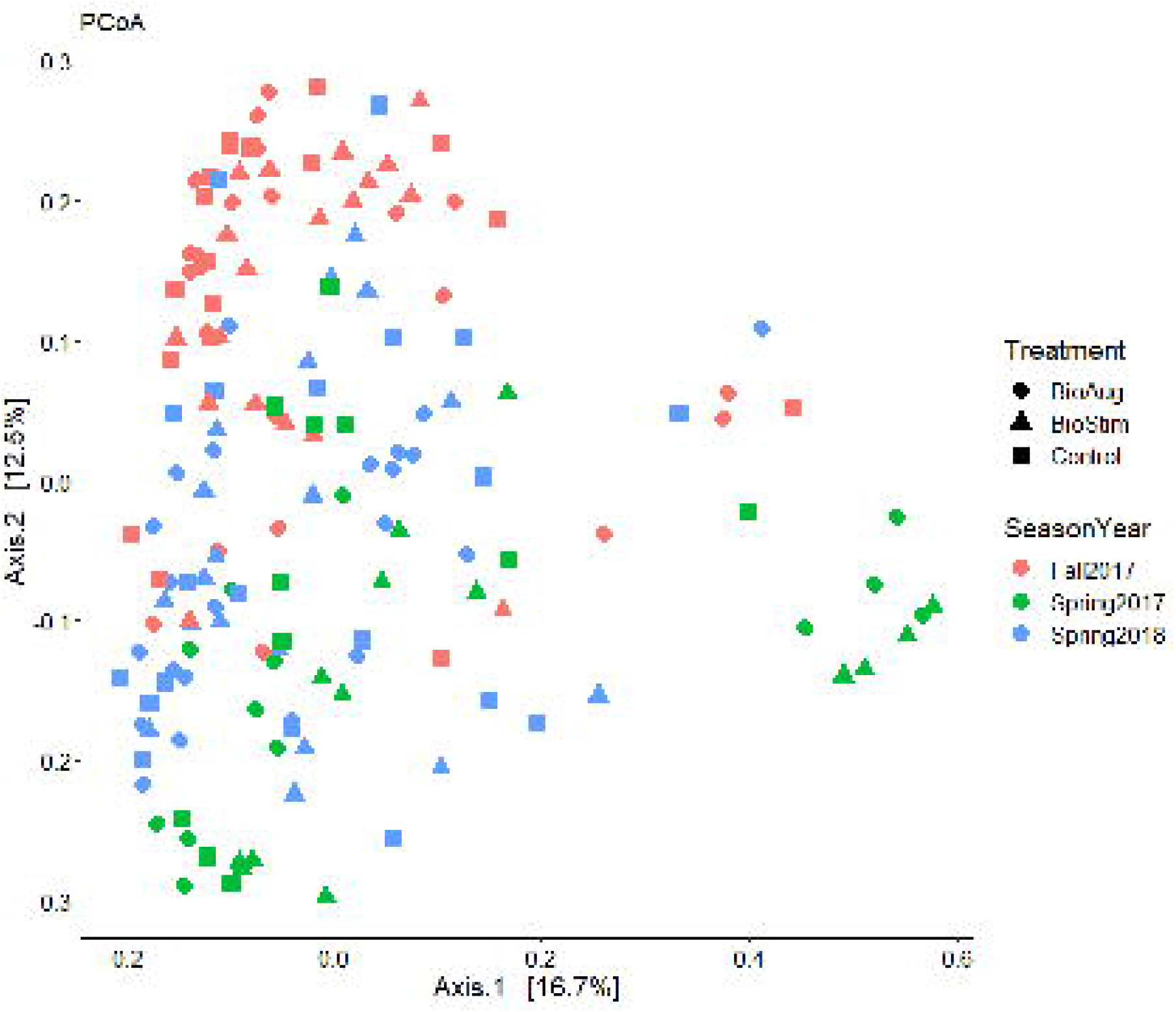
Principal coordinates analysis (PCoA) plot showing the Bary-Curtis dissimilarities among microbial communities in woodchip bioreactors. Each community is labeled with the treatment (the inoculation of cold-adapted denitrifiers [i.e., bioaugmentation], the addition of acetate [i.e., biostimulation], and control) and season (Spring 2017, Fall 2017, and Spring 2018).

Constrained analysis of principal coordinates (CAP) was used to analyze the relationship between the environmental variables, the relative abundance of N cycle-related genes, and the composition of microbiomes measured by the 16S rRNA gene amplicon sequencing (Fig. 4). Based on the adjusted r^2^ value of the CAP model, DO, ORP, temperature, conductivity, port number (location in the bioreactors), abundance of bacterial and archaeal 16S rRNA genes, relative abundance of *amoA, napA, nirK, nirS*, and *hdh* genes, and the relative abundance of inoculants (strain BE2.4 and strain WB94) were identified as the factors that can significantly (*p* <0.05) explain the difference in the microbiomes (Fig. 4). The CAP model can explain 33.5% of the variation within the microbiomes.

**Figure 4.**
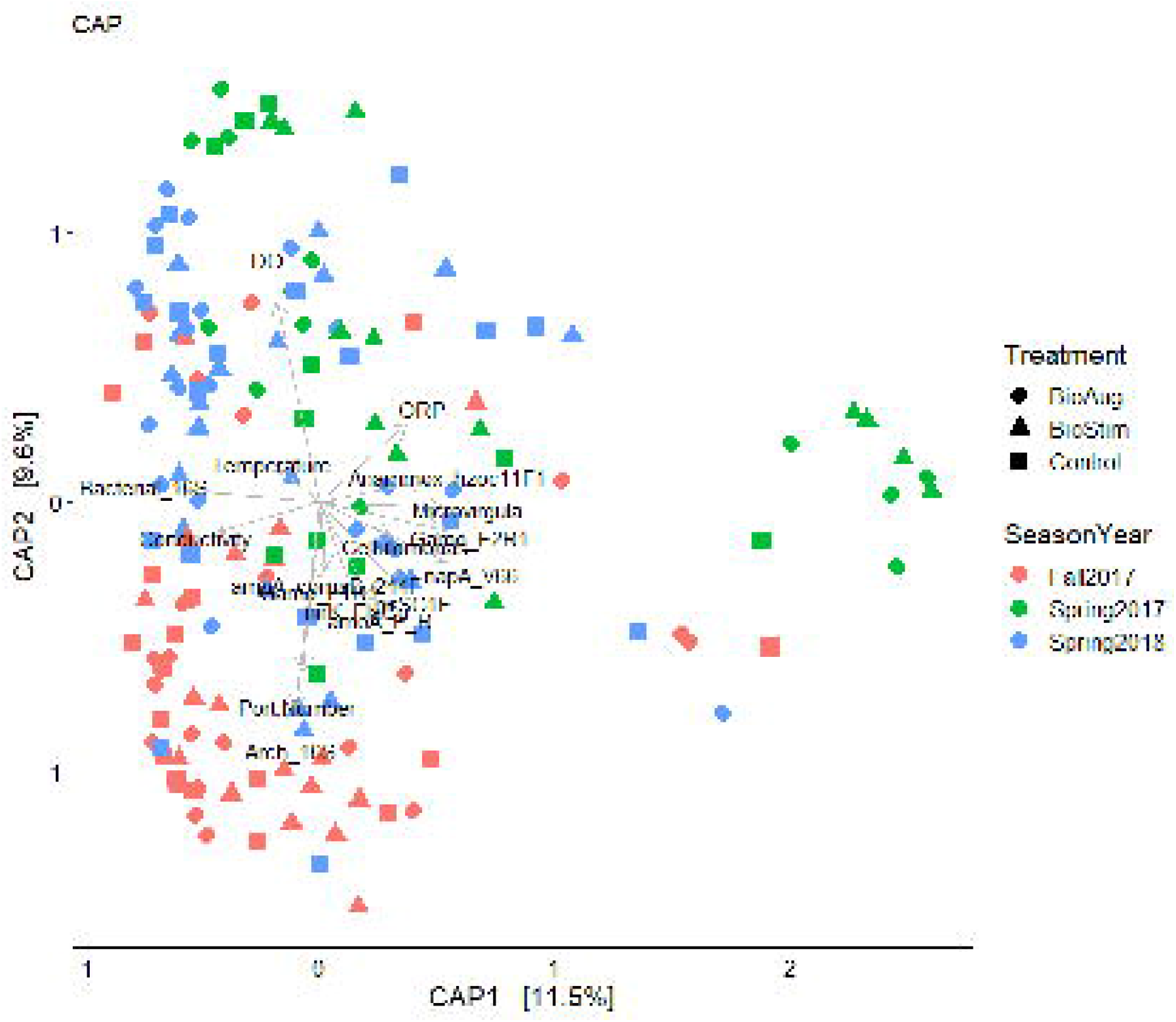
Constrained analysis of principal coordinates (CAP) biplot showing the association between the microbial communities (dots) and the N cycle gene abundance and environmental parameters (arrows). Microbial communities were analyzed using Bray-Curtis distance matrices. All environmental variables shown in the plots had significant effects (*p* <0.05) on the microbial community patterns based on PERMANOVA. Each community is labeled with the treatment (the inoculation of cold-adapted denitrifiers [i.e., bioaugmentation], the addition of acetate [i.e., biostimulation], and control) and season (Spring 2017, Fall 2017, and Spring 2018).

### Discussion

While N removal by WBR has been studied for more than 10 years, most studies focus on engineering and water chemistry aspects of WBR. Relatively little is known about the microbiological ecology in WBR, although microbes play a key role in WBR (Zhao et al., 2018; Aalto et al., 2020; Hartfiel et al., 2022). This study reports on temporal dynamics in the microbiome and the N cycle gene abundance in field-scale replicated WBR with different treatments (biostimulation, bioaugmentation, and control).

Our NiCE chip data suggests that there are denitrification hotspots in WBRs. The distribution of nitrogen cycle genes inside WBRs is rather uneven, and both nitrification and denitrification marker genes were found at elevated concentrations near the inlet of the WBRs. This may be related to the higher substrate (N and C) concentrations (Feyereisen et al., 2022). Dissolved oxygen concentration was also higher near the inlet of the WBRs, but it was rapidly consumed most likely by aerobic heterotrophic microbes (Feyereisen et al., 2022). Some microbes can also perform denitrification in aerobic conditions (Ji et al., 2015; Jang et al., 2018; Anderson et al., 2020). As a result, the region near the inlets of WBRs may be most suitable for nitrification and denitrification.

One of the advantages of the NiCE chip analysis is the ability to analyze multiple N cycle genes simultaneously. While the abundance of some of the denitrification maker genes (e.g., *nirS, nirK, nosZ*) has been analyzed in denitrifying bioreactors (Warneke et al., 2011; Schaefer et al., 2022), the intensive labor and cost required to perform multiple qPCR runs limits the number of N cycle genes being measured, and therefore, the abundance of other N cycle genes such as for nitrification and anammox have not been analyzed. Our NiCE chip results show that nitrifiers and anammox bacteria were also present in the WBRs, although ammonium concentration was rather low (<0.21 mg-N/L) and the temperature was low (Feyereisen et al., 2022). At present, it is unclear if nitrifiers and anammox bacteria are active in WBRs. RNA-based analysis may be needed to clarify if they are active in WBRs. Depending on the reactor operating conditions (e.g., high C/N biostimulation), DNRA bacteria can occur in WBRs (Anderson et al., 2020); therefore, under such conditions, nitrifiers and anammox bacteria may proliferate. The occurrence of nitrification and anammox in WBRs remains a future task.

We observed a significant shift in microbiome structures throughout the experimental period. The microbiome of samples from the same season (e.g., Spring) was more similar to each other than those of the samples from the same year (e.g., 2017), suggesting that there is a seasonal pattern in microbiome structures in WBRs. Our CAP model indicates that DO and temperature are among the variables significantly (*p* <0.05) associated with the patterns in the microbiome structure. While the DO concentration in woodchip bioreactors is associated with the sampling location, there is a seasonal change in temperature (Feyereisen et al., 2022). The average temperature of the Fall 2017 sampling campaign was higher than that of Spring 2017 and Spring 2018. Thus temperature could be a contributing factor to our results. Our findings contrast the previous study reporting that microbiomes in the samples collected from the same year were more comparable to each other than those from the same season in a two-year study in Iowa (Schaefer et al., 2022). This may be due, in part, to different WBR configurations, climate regions (i.e., colder winter in Minnesota than in Iowa), and the limited length of time in our study (i.e., no samples were collected in Fall 2016 and Fall 2018).

Although treatment had a significant impact on the microbiome structure, its influence was rather limited. Probe-based qPCR results of inoculants also suggest that the abundance of inoculated strains decreased within 4-6 weeks after the inoculation (Feyereisen et al., 2022). Mardani et al. (2020) also found that some of the inoculated microbes passed through the bioreactor instead of colonizing on woodchips and establishing robust populations. For the prolonged effect of bioaugmentation, inoculation techniques should be refined, for example, by encapsulating bacteria (Wang et al. 2020).

In this study, all the woodchip samples were collected below the water table. Due to the study design the flow rate and water table level of WBR were relatively constant. Previous studies suggest that microbiomes are distinct between the saturated and unsaturated portions of WBR (Porter et al., 2015, Hathaway et al., 2017); therefore, if we had also collected samples from the unsaturated zone the WBR, we may have observed greater differences in the WBR microbiomes.

Our GLM modeling provided some insights into the relationship between nitrogen removal rate and the abundance of N cycle-related genes. The NiCE chip data combined with the TaqMan probe strain-specific qPCR data showed that the abundance of inoculant strain BE2.4 had a positive relationship with the abundance of denitrification genes *nosZ* and *norB* in WBRs. Since both *nosZ* and *norB* have positive relationships with the NRR of WBRs based on the GLM, this suggests that bioaugmentation can increase the abundance of denitrification genes and improve the performance of WBRs.

In this study, we created multiple models to examine the relationship between the abundance of N cycle-related genes, environmental variables, the concentration of inoculants in the WBRs, and the microbiome structure measured by the 16S rRNA gene amplicon sequencing. However, some of the variables were strongly correlated with each other (e.g., port location and DO), which resulted in a high VIF value in the CAP analysis. Similar to this study, confounding factors can make it difficult to distinguish the interaction between WBR microbiome and environmental variables (Schaefer et al., 2022). To more clearly understand the relationships between the environmental variables and marker gene abundance, network analysis would be useful (Faust & Raes, 2012; Matchado et al., 2021). This should be tested in the future.

In conclusion, our NiCE chip and 16S rRNA gene amplicon sequencing results revealed temporal and spatial dynamics in microbiomes in WBR. The WBR microbiomes are likely not uniform, indicating there are denitrification hotspots in WBR. Based on our NiCE results, in addition to denitrification, other N cycle reactions may occur in WBR, such as nitrification and anammox, depending on the WBR operating conditions. The WBR microbiomes change by season, most likely in response to temperature and other environmental conditions (e.g., DO). Treatment (i.e., biostimulation and bioaugmentation) also influenced the overall WBR microbiome structures, although the effect was rather small. Inoculation of cold-adapted denitrifiers increased the denitrification functional genes, which can influence the NRR of WBRs.

Further research is needed, including to identify the key microbial species in nitrate removal in WBRs and analyze the environmental factors influencing their behavior. The WBR microbiomes are complex and only a small portion of the microbial community likely play a role in nitrate removal. Therefore, RNA-based analysis may be more useful to identify metabolically active microbes and directly link microbial activity to

N removal in WBR. Longer-term monitoring of microbiomes is also important to analyze temporal dynamics in the field-scale WBRs and the persistence of the inoculated bacteria.

## MATERIALS AND METHODS

### Woodchip sample collection

Woodchip samples were collected from six replicated WBR located in Willmar, MN. The WBR design is described in detail (Feyereisen et al., 2022). Briefly, each bioreactor was ≈ 11.6-m long and 1.7-m wide. The bioreactor beds received agricultural subsurface drainage from adjacent crop fields. Typical influent concentrations for nitrate-N and total organic carbon (TOC) ranged from 14 to 20 mg-N/L and 4.3 to 7.8 mg-C/L, respectively. Agricultural drainage was pumped into and stored in a 11.4 m^3^-L tank then fed to each WBR bed through polyvinyl chloride (PVC) piping. The water flow rate was controlled by manual valves and was relatively constant. Paddlewheel flow sensors were used to measure the flow rate. Five 15-cm diameter PVC pipes (“ports”) were installed vertically along the length of each bed that were used for water quality monitoring and woodchip sampling (Fig. S4a). Into each port was inserted a woodchip basket, which was filled with about 30 woodchip bags, each approximately 8-cm-diameter mesh bag containing about 100 g of woodchip (Fig. S4b).

The six WBR beds were divided into three treatments: biostimulation (beds #4 and #6), bioaugmentation (beds #3 and #7), and control (beds #5 and #8). For biostimulation, labile carbon (sodium acetate trihydrate) was injected via peristaltic pump to achieve C/N ratios ranging from 0.15 to 2.5 (Feyereisen et al., 2022). For bioaugmentation, two cold-adapted denitrifiers isolated from WBR, *Cellulomonas cellasea* strain WB94 (Jang et al., 2019) and *Microvirgula aerodenitrificans* strain BE2.4 (Anderson et al., 2020), were inoculated to the WBR. Bed flow rate was reduced in the bioaugmentation treatment beds and left low for one day after inoculations, except for one week after 17 October 2017, to keep the inoculant stay longer in the bed and improve the effectiveness of the bioaugmentation treatment. Control beds received neither acetate nor bacteria.

Woodchip samples were collected twice in Spring (May-June) 2017 and every other week in Fall 2017 (October-November) and Spring 2018 (a total of eight sampling dates). For microbiome analysis, one woodchip bag was collected from each of the sampling ports #1, #2, #3, #4, and #5 (Fig. S4a) on the two sampling dates during the Spring 2017 field campaign. For the Fall 2017 and Spring 2018 sampling campaigns, the woodchip bags were collected from sampling ports #2, #3, #4, and #5. The woodchip bags collected were immediately stored in a cooler with ice packs, transported to the laboratory on the same day, and stored in a −20°C freezer until used. The list of samples collected and analyzed is shown in Table S1.

Various physicochemical properties of the WBR water, including oxidation-reduction potential (ORP), temperature, conductivity, dissolved oxygen (DO) concentration, and pH were measured from the same port by pumping water from 5 cm from the bioreactor bottom with a peristaltic pump into a 0.75-L vessel, allowing the water to rise above a submerged multi-parameter sonde. Water samples were also collected to analyze nitrate and dissolved organic carbon (DOC) concentrations as described by Feyereisen et al. (2022). The NRR of each WBR bed was calculated and normalized to gram of NO_3_-N removed per cubic meter of WBR bed per day (g N m^-3^ d^-1^) (Feyereisen et al., 2022). In addition, the abundance of inoculated strains, *Microvirgula aerodenitrificans* strain BE2.4 and *Cellulomonas cellasea* strain WB94, was measured by strain-specific qPCR (Feyereisen et al., 2022).

### DNA extraction of woodchip samples

The woodchip bags taken out from the −20°C freezer were left at room temperature for 40 min to partially thaw. Then, 25 g of woodchips were placed into a 160 mL wide-mouth milk dilution bottle (Corning) containing 100 mL of phosphate-buffered saline solution (pH 7.5) with 0.1% gelatin (PBS-gelatin) and 25 g of 5-mm-diam. glass beads. The milk bottles were placed on a shaker and shaken horizontally for 30 minutes to release bacterial cells from the woodchips. The bacteria suspension in PBS-gelatin buffer was then transferred to a 50 mL centrifuge tube (Thermo Scientific) and centrifuged at 10,000 rpm (11,953 ×*g*) for 15 min at 4°C. After the centrifugation, the supernatant was discarded, and this process was repeated until all the PBS-gelatin buffer from the milk bottle was transferred and centrifuged. The bacterial pellet was weighed and stored in a 2-ml centrifuge tube at −80°C until further processed. A total of 170 woodchip samples were collected, of which 169 samples were used for DNA extraction. One sample was discarded due to mislabeling.

The PowerSoil DNA extraction kit (Qiagen) was used to extract DNA from the bacterial pellet washed off from the woodchips. The extraction was done using the QIAcube Connect automated system (Qiagen) following the manufacturer’s protocol, with the exception that 0.5 g of the bacterial pellet was used for the extraction instead of 0.25 g of soil. The DNA elution was diluted 10-fold and stored in the −80°C freezer. The quality of DNA was verified by qPCR targeting the 16S rRNA gene as described by Jang et al. (2019).

### Nitrogen cycle evaluation (NiCE) chip

The DNA samples (n=169) were subjected to the nitrogen cycle evaluation (NiCE) chip (Oshiki et al., 2018, Jang et al., 2022) to quantify various nitrogen cycle-associated genes, including those for nitrification (*amoA, hao*, and *nxrB*), denitrification (*napA, narG, nirK, nirS, norB*, and *nosZ* clade I & II), dissimilatory nitrate reduction to ammonium (DNRA; *nrfA*), anaerobic ammonium oxidation (anammox; *hdh* and *hzs*), and nitrogen fixation (*nifH*). A list of the assays used is shown in Table S2. Synthetic gBlock DNA fragments (Integrated DNA Technologies) containing the target gene fragments were used to create the standards for qPCR (Jang et al., 2022). The gBlock gene fragments were pooled together and serially diluted (3.34 × 10^7^ to 3.34 *×* 10^0^ copies/μl) to generate standard curves.

The NiCE chip was run using the SmartChip MyDesign chip (Takara Bio) and the SmartChip Real-Time PCR system (Takara Bio). The SmartChip MyDesign chip contains 5,184 nanowells, into which all possible combinations of 36 assays and 144 samples (127 woodchip samples, 16 standards, and one no-template control) were loaded by the SmartChip MultiSample NanoDispenser (Takara Bio). In brief, 50 nl each of the DNA samples mixed with 1X SmartChip TB Green Gene Expression Master Mix (Takara Bio) were first loaded onto the SmartChip MyDesign Chip. Then 50 nl of primer pair solution mixed with 1X SmartChip TB Green Gene Expression Master Mix (Takara Bio) were loaded onto the SmartChip. The final volume in each nanowell of the MyDesign chip was 100 nl, and the final primer pair concentration in each nanowell was 500 nM. The qPCR reaction was done under the following thermal condition: 95°C for 3 min followed by 40 cycles at 95°C for 30 sec, 50°C for 30 sec, and 72°C for 30 sec. Melting curve analysis was performed from 50°C to 97°C with a 0.4°C/step temperature gradient.

The threshold cycle (Ct) values were determined by the SmartChip Real-Time PCR software (Takara Bio). Standard curves were generated by plotting the Ct values vs. the concentration (copies/μl) of standard DNA. Standard curves with at least three data points and an r^2^ value of >0.95 were considered valid. Target gene concentrations in the woodchip samples were determined based on their Ct values by using the standard curves (Ishii et al. 2013). For samples that showed below the limit of quantification (LOQ), LOQ/2 were assigned as recommended by Hites (2019). The nitrogen cycle-associated gene concentrations were normalized by dividing them by the 16S rRNA gene concentration. Two samples had low 16S rRNA gene concentration and did not produce positives in more than half of the NiCE chip assays, and therefore, were excluded from the downstream analysis.

### 16S rRNA gene amplicon sequencing

The same DNA samples (n=169) were also used for the 16S rRNA gene amplicon sequencing to analyze bacterial/archaeal communities in the WBR. The V4 region of the 16S rRNA gene was amplified, purified, and used to prepare sequencing libraries according to Gohl et al. (2016). The 300-bp paired-end sequencing was done using the MiSeq platform (Illumina) with V3 chemistry. The 16S rRNA gene amplification, library preparation, and sequencing were done at the University of Minnesota Genomics Center (UMGC).

The sequence reads were quality-filtered, assembled, and trimmed using the NINJA-SHI7 (Al-Ghalith et al., 2018). The output file generated by NINJA-SHI7 were then processed through the NINJA-OPS pipeline (Al-Ghalith et al., 2016). The sequences were clustered to operational taxonomic units (OTUs) at 97% similarity. Taxonomic information was assigned using NINJA-OPS with the Greengene database ver. 13_8 (McDonald et al., 2012). The biom file created by the NINJA-OPS pipeline was used for statistical analysis.

### Statistical analysis

Microbiome structures were analyzed by using RStudio version 4.1.0 with *vegan* (Oksanen et al., 2019) and *physeq* packages (McMurdie & Holmes, 2013). The numbers of sequences were normalized by rarefaction (Jang et al., 2022). Permutational multivariate analysis of variance (PERMANOVA) was used to examine the differences in the microbiome structures. Principal coordinates analysis (PCoA) with Bray-Curtis distance matrices were used to visualize the dissimilarities in microbiome structures of woodchip samples. Canonical analysis of principal coordinates (CAP) was used to identify the environmental variables and nitrogen cycle genes associated with the microbiome structures. Variance inflation factors (VIF) were calculated to examine the correlation between environmental variables and nitrogen cycle gene abundance. Variables with a VIF of >10 were removed from the CAP model (Jang et al., 2022).

Paired t-tests were conducted for the average of the twelve denitrification genes in samples collected from Port #2 and Port #5 using Excel. Spearman’s correlations between environmental variables, N-cycle associated genes, and abundance of the inoculated bacteria were analyzed and visualized by using the *corrplot* package (Wei & Simko, 2021). Generalized linear model (GLM) was used to identify nitrogen cycle genes that can predict the changes in weekly NRR of WBRs.

## Data availability

The 16S rRNA gene sequences generated in this study were submitted to the NCBI Sequence Read Archive and are available under BioProject number PRJNA887466.

## ACKNOWLEDGMENTS

This study was supported by MnDRIVE Environment Initiative, Minnesota Department of Agriculture (Project No. 108837), Minnesota Agricultural Water Resource Center/Discovery Farms Minnesota, University of Minnesota Department of Soil, Water, and Climate, and USDA-ARS.

We also thank Emily Anderson, Ed Dorsey, Scott Schumacher, Todd Schumacher, Chan Lan Chun, Jeonghwan Jang, and Andry Ranaivoson, and a host of staff and students.

## DISCLAIMER

Mention of trade names or commercial products in this publication is solely for the purpose of providing specific information and does not imply recommendation or endorsement by the U.S. Department of Agriculture. USDA is an equal opportunity provider and employer.

