## Supplementary material for "Impacts of Biostimulation and Bioaugmentation on Woodchip Bioreactor Microbiomes": Fig. S1, Fig. S2, Fig. S3, Fig. S4, Table S1, Table S2, Table S3

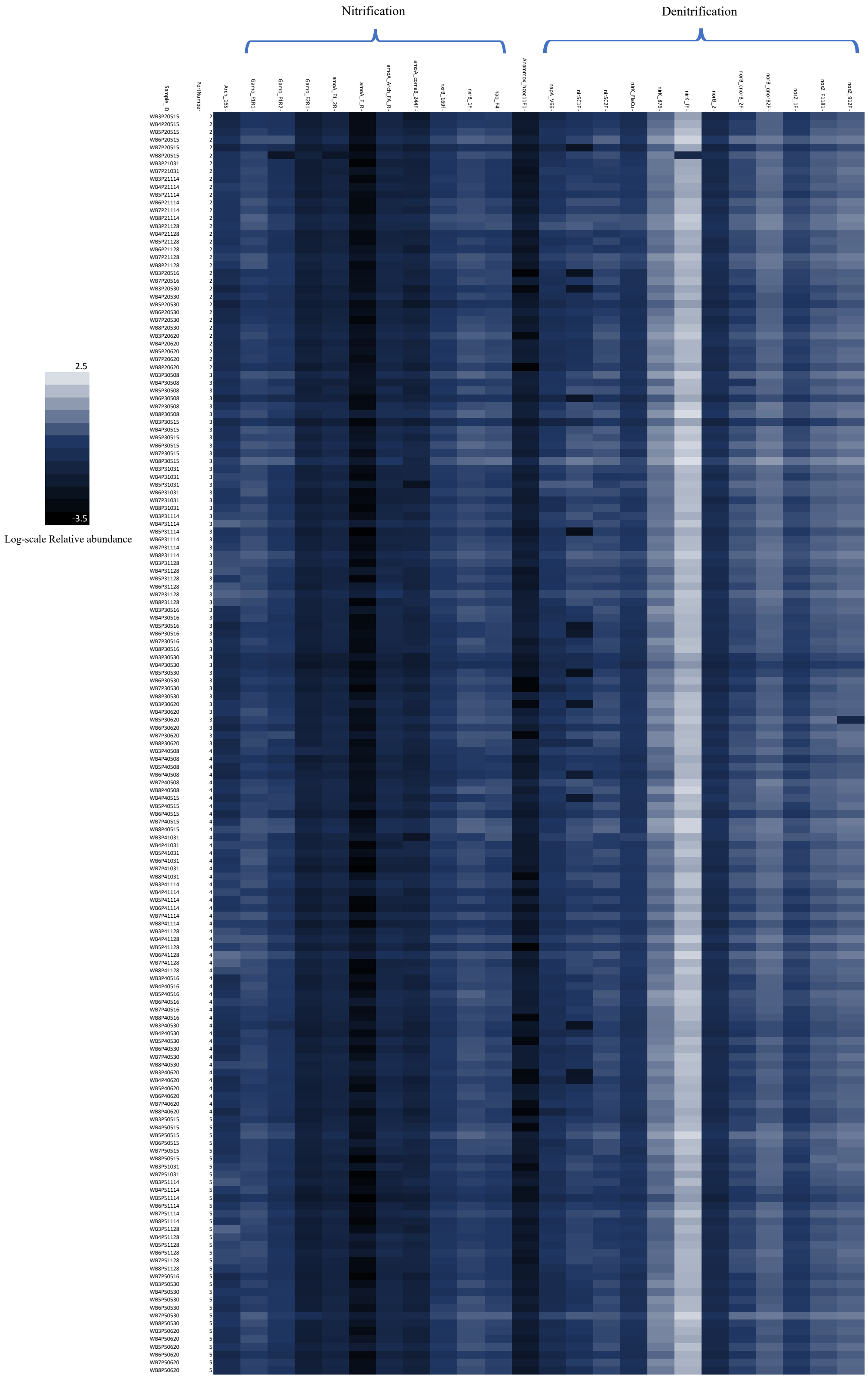


**Figure S1.** Heatmap showing the abundances of nitrification and denitrification genes in the woodchip bioreactor samples as measured by the Nitrogen Cycle Evaluation (NiCE) chip. See Table S1 for sample ID.

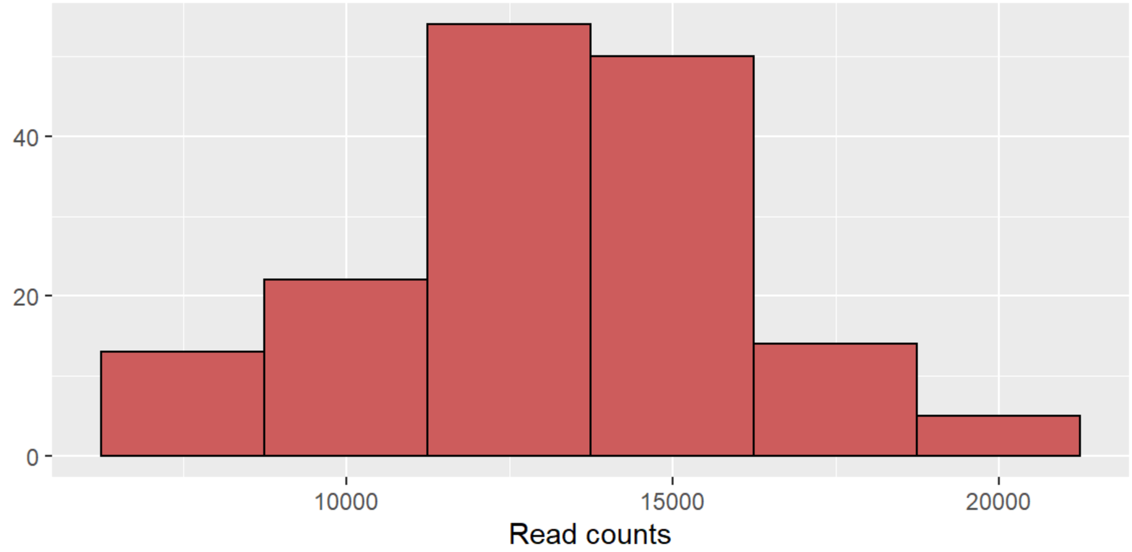


**Figure S2.** Distribution of sample sequencing depth (n = 158; mean = 13,153 reads; median = 13,275 reads)

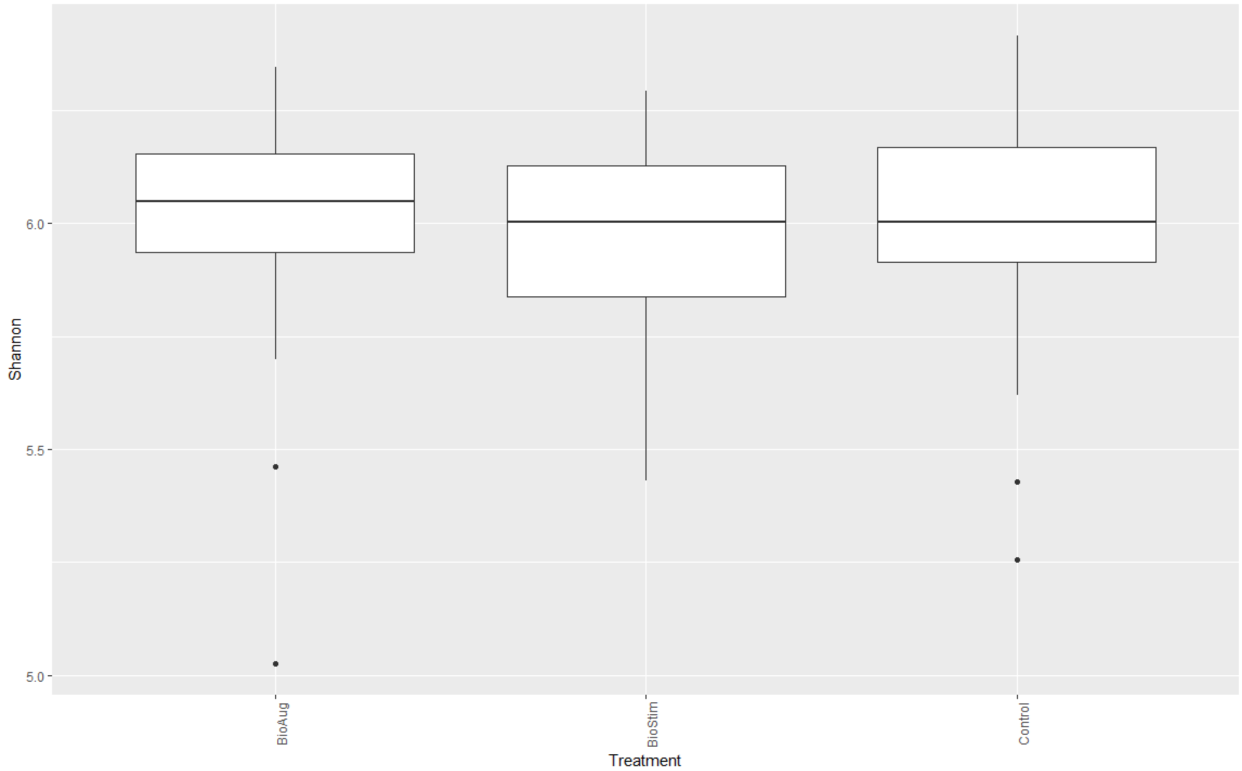


**Figure S3.** Shannon index of samples collected from different treatment groups.

(a)

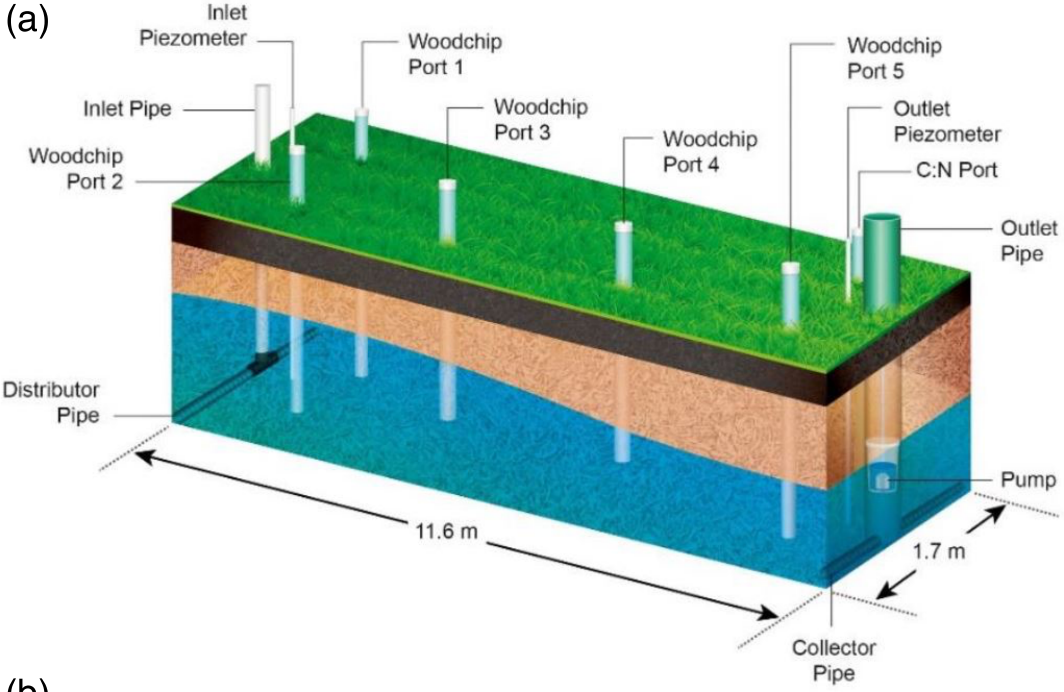


(b)

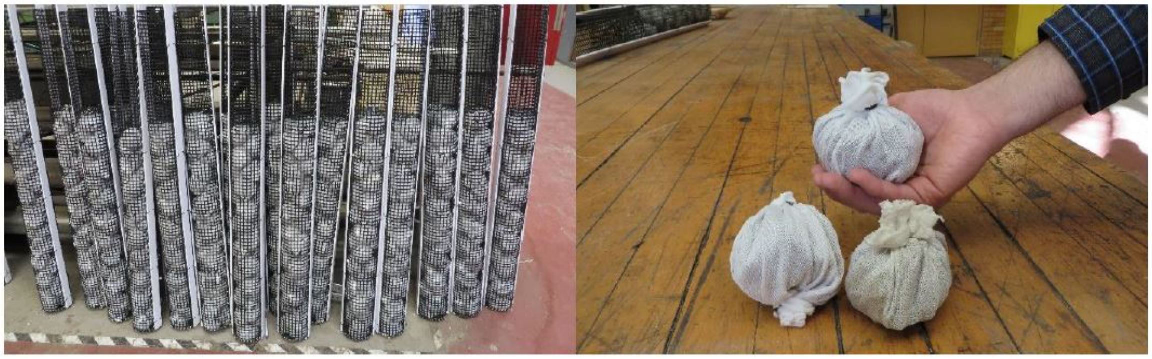

**Figure S4.** Diagrams of woodchip bioreactors. (a) Design of the woodchip bioreactor bed with ports for woodchip sampling. (b) Photo of woodchip baskets inside the sampling ports and “woodchip balls” placed in the sampling ports. Adapted from Feyereisen et al. (2018).

**Table S1.** Sample description.

| Sample ID | Port ID | Reactor No. | Port No. | Treatment | Date |
|---|---|---|---|---|---|
| WB3P30508 | WB3P3 | 3 | 3 | Bio-Aug | 2017-05-08 |
| WB3P40508 | WB3P4 | 3 | 4 | Bio-Aug | 2017-05-08 |
| WB7P30508 | WB7P3 | 7 | 3 | Bio-Aug | 2017-05-08 |
| WB7P40508 | WB7P4 | 7 | 4 | Bio-Aug | 2017-05-08 |
| WB4P30508 | WB4P3 | 4 | 3 | Bio-Stim | 2017-05-08 |
| WB4P40508 | WB4P4 | 4 | 4 | Bio-Stim | 2017-05-08 |
| WB5P30508 | WB5P3 | 5 | 3 | Control | 2017-05-08 |
| WB5P40508 | WB5P4 | 5 | 4 | Control | 2017-05-08 |
| WB6P30508 | WB6P3 | 6 | 3 | Bio-Stim | 2017-05-08 |
| WB6P40508 | WB6P4 | 6 | 4 | Bio-Stim | 2017-05-08 |
| WB8P30508 | WB8P3 | 8 | 3 | Control | 2017-05-08 |
| WB8P40508 | WB8P4 | 8 | 4 | Control | 2017-05-08 |
| WB3P10515 | WB3P1 | 3 | 1 | Bio-Aug | 2017-05-15 |
| WB3P20515 | WB3P2 | 3 | 2 | Bio-Aug | 2017-05-15 |
| WB3P30515 | WB3P3 | 3 | 3 | Bio-Aug | 2017-05-15 |
| WB3P40515 | WB3P4 | 3 | 4 | Bio-Aug | 2017-05-15 |
| WB3P50515 | WB3P5 | 3 | 5 | Bio-Aug | 2017-05-15 |
| WB7P10515 | WB7P1 | 7 | 1 | Bio-Aug | 2017-05-15 |
| WB7P20515 | WB7P2 | 7 | 2 | Bio-Aug | 2017-05-15 |
| WB7P30515 | WB7P3 | 7 | 3 | Bio-Aug | 2017-05-15 |
| WB7P40515 | WB7P4 | 7 | 4 | Bio-Aug | 2017-05-15 |
| WB7P50515 | WB7P5 | 7 | 5 | Bio-Aug | 2017-05-15 |
| WB4P10515 | WB4P1 | 4 | 1 | Bio-Stim | 2017-05-15 |
| WB4P20515 | WB4P2 | 4 | 2 | Bio-Stim | 2017-05-15 |
| WB4P30515 | WB4P3 | 4 | 3 | Bio-Stim | 2017-05-15 |
| WB4P40515 | WB4P4 | 4 | 4 | Bio-Stim | 2017-05-15 |
| WB4P50515 | WB4P5 | 4 | 5 | Bio-Stim | 2017-05-15 |
| WB5P10515 | WB5P1 | 5 | 1 | Control | 2017-05-15 |
| WB5P20515 | WB5P2 | 5 | 2 | Control | 2017-05-15 |
| WB5P30515 | WB5P3 | 5 | 3 | Control | 2017-05-15 |
| WB5P40515 | WB5P4 | 5 | 4 | Control | 2017-05-15 |
| WB5P50515 | WB5P5 | 5 | 5 | Control | 2017-05-15 |
| WB6P10515 | WB6P1 | 6 | 1 | Bio-Stim | 2017-05-15 |
| WB6P20515 | WB6P2 | 6 | 2 | Bio-Stim | 2017-05-15 |
| WB6P30515 | WB6P3 | 6 | 3 | Bio-Stim | 2017-05-15 |
| WB6P40515 | WB6P4 | 6 | 4 | Bio-Stim | 2017-05-15 |

| | | | | | |
|---|---|---|---|---|---|
| WB6P50515 | WB6P5 | 6 | 5 | Bio-Stim | 2017-05-15 |
| WB8P10515 | WB8P1 | 8 | 1 | Control | 2017-05-15 |
| WB8P20515 | WB8P2 | 8 | 2 | Control | 2017-05-15 |
| WB8P30515 | WB8P3 | 8 | 3 | Control | 2017-05-15 |
| WB8P40515 | WB8P4 | 8 | 4 | Control | 2017-05-15 |
| WB8P50515 | WB8P5 | 8 | 5 | Control | 2017-05-15 |
| WB3P21031 | WB3P2 | 3 | 2 | Bio-Aug | 2017-10-31 |
| WB3P31031 | WB3P3 | 3 | 3 | Bio-Aug | 2017-10-31 |
| WB3P41031 | WB3P4 | 3 | 4 | Bio-Aug | 2017-10-31 |
| WB3P51031 | WB3P5 | 3 | 5 | Bio-Aug | 2017-10-31 |
| WB7P21031 | WB7P2 | 7 | 2 | Bio-Aug | 2017-10-31 |
| WB7P31031 | WB7P3 | 7 | 3 | Bio-Aug | 2017-10-31 |
| WB7P41031 | WB7P4 | 7 | 4 | Bio-Aug | 2017-10-31 |
| WB7P51031 | WB7P5 | 7 | 5 | Bio-Aug | 2017-10-31 |
| WB4P31031 | WB4P3 | 4 | 3 | Bio-Stim | 2017-10-31 |
| WB4P41031 | WB4P4 | 4 | 4 | Bio-Stim | 2017-10-31 |
| WB5P31031 | WB5P3 | 5 | 3 | Control | 2017-10-31 |
| WB5P41031 | WB5P4 | 5 | 4 | Control | 2017-10-31 |
| WB6P31031 | WB6P3 | 6 | 3 | Bio-Stim | 2017-10-31 |
| WB6P41031 | WB6P4 | 6 | 4 | Bio-Stim | 2017-10-31 |
| WB8P31031 | WB8P3 | 8 | 3 | Control | 2017-10-31 |
| WB8P41031 | WB8P4 | 8 | 4 | Control | 2017-10-31 |
| WB3P21114 | WB3P2 | 3 | 2 | Bio-Aug | 2017-11-14 |
| WB3P31114 | WB3P3 | 3 | 3 | Bio-Aug | 2017-11-14 |
| WB3P41114 | WB3P4 | 3 | 4 | Bio-Aug | 2017-11-14 |
| WB3P51114 | WB3P5 | 3 | 5 | Bio-Aug | 2017-11-14 |
| WB7P21114 | WB7P2 | 7 | 2 | Bio-Aug | 2017-11-14 |
| WB7P31114 | WB7P3 | 7 | 3 | Bio-Aug | 2017-11-14 |
| WB7P41114 | WB7P4 | 7 | 4 | Bio-Aug | 2017-11-14 |
| WB7P51114 | WB7P5 | 7 | 5 | Bio-Aug | 2017-11-14 |
| WB4P21114 | WB4P2 | 4 | 2 | Bio-Stim | 2017-11-14 |
| WB4P31114 | WB4P3 | 4 | 3 | Bio-Stim | 2017-11-14 |
| WB4P41114 | WB4P4 | 4 | 4 | Bio-Stim | 2017-11-14 |
| WB4P51114 | WB4P5 | 4 | 5 | Bio-Stim | 2017-11-14 |
| WB5P21114 | WB5P2 | 5 | 2 | Control | 2017-11-14 |
| WB5P31114 | WB5P3 | 5 | 3 | Control | 2017-11-14 |
| WB5P41114 | WB5P4 | 5 | 4 | Control | 2017-11-14 |
| WB5P51114 | WB5P5 | 5 | 5 | Control | 2017-11-14 |
| WB6P21114 | WB6P2 | 6 | 2 | Bio-Stim | 2017-11-14 |

| | | | | | |
|---|---|---|---|---|---|
| WB6P31114 | WB6P3 | 6 | 3 | Bio-Stim | 2017-11-14 |
| WB6P41114 | WB6P4 | 6 | 4 | Bio-Stim | 2017-11-14 |
| WB6P51114 | WB6P5 | 6 | 5 | Bio-Stim | 2017-11-14 |
| WB8P21114 | WB8P2 | 8 | 2 | Control | 2017-11-14 |
| WB8P31114 | WB8P3 | 8 | 3 | Control | 2017-11-14 |
| WB8P41114 | WB8P4 | 8 | 4 | Control | 2017-11-14 |
| WB8P51114 | WB8P5 | 8 | 5 | Control | 2017-11-14 |
| WB3P21128 | WB3P2 | 3 | 2 | Bio-Aug | 2017-11-28 |
| WB3P31128 | WB3P3 | 3 | 3 | Bio-Aug | 2017-11-28 |
| WB3P41128 | WB3P4 | 3 | 4 | Bio-Aug | 2017-11-28 |
| WB3P51128 | WB3P5 | 3 | 5 | Bio-Aug | 2017-11-28 |
| WB7P21128 | WB7P2 | 7 | 2 | Bio-Aug | 2017-11-28 |
| WB7P31128 | WB7P3 | 7 | 3 | Bio-Aug | 2017-11-28 |
| WB7P41128 | WB7P4 | 7 | 4 | Bio-Aug | 2017-11-28 |
| WB7P51128 | WB7P5 | 7 | 5 | Bio-Aug | 2017-11-28 |
| WB4P21128 | WB4P2 | 4 | 2 | Bio-Stim | 2017-11-28 |
| WB4P31128 | WB4P3 | 4 | 3 | Bio-Stim | 2017-11-28 |
| WB4P41128 | WB4P4 | 4 | 4 | Bio-Stim | 2017-11-28 |
| WB4P51128 | WB4P5 | 4 | 5 | Bio-Stim | 2017-11-28 |
| WB5P21128 | WB5P2 | 5 | 2 | Control | 2017-11-28 |
| WB5P31128 | WB5P3 | 5 | 3 | Control | 2017-11-28 |
| WB5P41128 | WB5P4 | 5 | 4 | Control | 2017-11-28 |
| WB5P51128 | WB5P5 | 5 | 5 | Control | 2017-11-28 |
| WB6P21128 | WB6P2 | 6 | 2 | Bio-Stim | 2017-11-28 |
| WB6P31128 | WB6P3 | 6 | 3 | Bio-Stim | 2017-11-28 |
| WB6P41128 | WB6P4 | 6 | 4 | Bio-Stim | 2017-11-28 |
| WB6P51128 | WB6P5 | 6 | 5 | Bio-Stim | 2017-11-28 |
| WB8P21128 | WB8P2 | 8 | 2 | Control | 2017-11-28 |
| WB8P31128 | WB8P3 | 8 | 3 | Control | 2017-11-28 |
| WB8P41128 | WB8P4 | 8 | 4 | Control | 2017-11-28 |
| WB8P51128 | WB8P5 | 8 | 5 | Control | 2017-11-28 |
| WB3P20516 | WB3P2 | 3 | 2 | Bio-Aug | 2018-05-16 |
| WB3P30516 | WB3P3 | 3 | 3 | Bio-Aug | 2018-05-16 |
| WB3P40516 | WB3P4 | 3 | 4 | Bio-Aug | 2018-05-16 |
| WB3P50516 | WB3P5 | 3 | 5 | Bio-Aug | 2018-05-16 |
| WB7P20516 | WB7P2 | 7 | 2 | Bio-Aug | 2018-05-16 |
| WB7P30516 | WB7P3 | 7 | 3 | Bio-Aug | 2018-05-16 |
| WB7P40516 | WB7P4 | 7 | 4 | Bio-Aug | 2018-05-16 |
| WB7P50516 | WB7P5 | 7 | 5 | Bio-Aug | 2018-05-16 |

| | | | | | |
|---|---|---|---|---|---|
| WB4P30516 | WB4P3 | 4 | 3 | Bio-Stim | 2018-05-16 |
| WB4P40516 | WB4P4 | 4 | 4 | Bio-Stim | 2018-05-16 |
| WB5P30516 | WB5P3 | 5 | 3 | Control | 2018-05-16 |
| WB5P40516 | WB5P4 | 5 | 4 | Control | 2018-05-16 |
| WB6P30516 | WB6P3 | 6 | 3 | Bio-Stim | 2018-05-16 |
| WB6P40516 | WB6P4 | 6 | 4 | Bio-Stim | 2018-05-16 |
| WB8P30516 | WB8P3 | 8 | 3 | Control | 2018-05-16 |
| WB8P40516 | WB8P4 | 8 | 4 | Control | 2018-05-16 |
| WB3P20530 | WB3P2 | 3 | 2 | Bio-Aug | 2018-05-30 |
| WB3P30530 | WB3P3 | 3 | 3 | Bio-Aug | 2018-05-30 |
| WB3P40530 | WB3P4 | 3 | 4 | Bio-Aug | 2018-05-30 |
| WB3P50530 | WB3P5 | 3 | 5 | Bio-Aug | 2018-05-30 |
| WB7P20530 | WB7P2 | 7 | 2 | Bio-Aug | 2018-05-30 |
| WB7P30530 | WB7P3 | 7 | 3 | Bio-Aug | 2018-05-30 |
| WB7P40530 | WB7P4 | 7 | 4 | Bio-Aug | 2018-05-30 |
| WB7P50530 | WB7P5 | 7 | 5 | Bio-Aug | 2018-05-30 |
| WB4P20530 | WB4P2 | 4 | 2 | Bio-Stim | 2018-05-30 |
| WB4P30530 | WB4P3 | 4 | 3 | Bio-Stim | 2018-05-30 |
| WB4P40530 | WB4P4 | 4 | 4 | Bio-Stim | 2018-05-30 |
| WB4P50530 | WB4P5 | 4 | 5 | Bio-Stim | 2018-05-30 |
| WB5P20530 | WB5P2 | 5 | 2 | Control | 2018-05-30 |
| WB5P30530 | WB5P3 | 5 | 3 | Control | 2018-05-30 |
| WB5P40530 | WB5P4 | 5 | 4 | Control | 2018-05-30 |
| WB5P50530 | WB5P5 | 5 | 5 | Control | 2018-05-30 |
| WB6P20530 | WB6P2 | 6 | 2 | Bio-Stim | 2018-05-30 |
| WB6P30530 | WB6P3 | 6 | 3 | Bio-Stim | 2018-05-30 |
| WB6P40530 | WB6P4 | 6 | 4 | Bio-Stim | 2018-05-30 |
| WB6P50530 | WB6P5 | 6 | 5 | Bio-Stim | 2018-05-30 |
| WB8P20530 | WB8P2 | 8 | 2 | Control | 2018-05-30 |
| WB8P30530 | WB8P3 | 8 | 3 | Control | 2018-05-30 |
| WB8P40530 | WB8P4 | 8 | 4 | Control | 2018-05-30 |
| WB8P50530 | WB8P5 | 8 | 5 | Control | 2018-05-30 |
| WB3P20620 | WB3P2 | 3 | 2 | Bio-Aug | 2018-06-20 |
| WB3P30620 | WB3P3 | 3 | 3 | Bio-Aug | 2018-06-20 |
| WB3P40620 | WB3P4 | 3 | 4 | Bio-Aug | 2018-06-20 |
| WB3P50620 | WB3P5 | 3 | 5 | Bio-Aug | 2018-06-20 |
| WB7P20620 | WB7P2 | 7 | 2 | Bio-Aug | 2018-06-20 |
| WB7P30620 | WB7P3 | 7 | 3 | Bio-Aug | 2018-06-20 |
| WB7P40620 | WB7P4 | 7 | 4 | Bio-Aug | 2018-06-20 |

| | | | | | |
|---|---|---|---|---|---|
| WB7P50620 | WB7P5 | 7 | 5 | Bio-Aug | 2018-06-20 |
| WB4P20620 | WB4P2 | 4 | 2 | Bio-Stim | 2018-06-20 |
| WB4P30620 | WB4P3 | 4 | 3 | Bio-Stim | 2018-06-20 |
| WB4P40620 | WB4P4 | 4 | 4 | Bio-Stim | 2018-06-20 |
| WB4P50620 | WB4P5 | 4 | 5 | Bio-Stim | 2018-06-20 |
| WB5P20620 | WB5P2 | 5 | 2 | Control | 2018-06-20 |
| WB5P30620 | WB5P3 | 5 | 3 | Control | 2018-06-20 |
| WB5P40620 | WB5P4 | 5 | 4 | Control | 2018-06-20 |
| WB5P50620 | WB5P5 | 5 | 5 | Control | 2018-06-20 |
| WB6P20620 | WB6P2 | 6 | 2 | Bio-Stim | 2018-06-20 |
| WB6P30620 | WB6P3 | 6 | 3 | Bio-Stim | 2018-06-20 |
| WB6P40620 | WB6P4 | 6 | 4 | Bio-Stim | 2018-06-20 |
| WB6P50620 | WB6P5 | 6 | 5 | Bio-Stim | 2018-06-20 |
| WB8P20620 | WB8P2 | 8 | 2 | Control | 2018-06-20 |
| WB8P30620 | WB8P3 | 8 | 3 | Control | 2018-06-20 |
| WB8P40620 | WB8P4 | 8 | 4 | Control | 2018-06-20 |
| WB8P50620 | WB8P5 | 8 | 5 | Control | 2018-06-20 |

**Table S2.** Primers used for the high-throughput N cycle quantification (Nitrogen Cycle Evaluation [NiCE] chip)

| Target | Assay ID | Primer name | Sequence (5' --> 3') | Reference |
|---|---|---|---|---|
| Total Archaea | Arch_16S | Archaea-F KO | CCCTAYGGGGYGCASCAGGC | Murakami et al. 2012 |
| | | Archaea-R KO | GCYCYCCCGCCAATTCMTTTA | |
| AOB (Gamma-proteobacteria) | Gamo_F1R1 | Gamo172 F1 | GGBGACTGGGAYTTCTGG | Oshiki et al. 2018 |
| | | Gamo172 F1_R1 | AAARCCCGAGAAGAAMGC | |
| | Gamo_F1R2 | Gamo172 F1 | GGBGACTGGGAYTTCTGG | |
| | | Gamo172 F1_R2 | AAAACCCGCAAAAAAGGC | |
| | Gamo_F2R1 | Gamo172 F2 | TGGGATTTCTGGATGGAC | |
| | | Gamo172 F2_R1 | TGATACGAACGCAGAGAA | |
| AOB (Beta-proteobacteria) | Bac_amoA | amoA_F1 | GGGGHTTYTACTGGTGGT | Rotthauwe et al. 1997; Oshiki et al. 2018 |
| | | amoA_2R | CCCCTCKGSAAAGCCTTCTTC | |
| AOB hao | hao | haoF4 | AYCTKCGCTCRATGGG | Schmid et al. 2008 |
| | | haoR2 | GGTTGGTYTTCTGKCCGG | |
| AOA | Arch_amoAF | Arch-amoAF | STAATGGTCTGGCTTAGACG | Francis et al. 2005 |
| | | Arch-amoAR | GCGGCCATCCATCTGTATGT | |
| | Arch_amoAFA | Arch-amoAFA | ACACCAGTTTGGYTACCWTCDGC | Beman et al. 2008 |
| | | Arch-amoAR | GCGGCCATCCATCTGTATGT | |
| | Arch_amoAFB* | Arch-amoAFB | CATCCRATGTGGATTCCATCDTG | |
| | | Arch-amoAR | GCGGCCATCCATCTGTATGT | |
| | Arch_amoA-for* | Arch-amoA-for | CTGAYTGGGCYTGGACATC | Wuchter et al. 2006 |
| | | Arch-amoA-rev | TTCTTCTTTGTTGCCCAGTA | |
| NOB (Nitrobacter) | nxrBF | NxrB 1F | ACGTGGAGACCAAGCCGGG | Vanparys et al. 2007 |
| | | NxrB 1R | CCGTGCTGTTGAYCTCGTTGA | |
| NOB (Nitrospira) | nxrB169f | nxrB169f | TACATGTGGTGGAACA | Pester et al. 2014 |
| | | nxrB638r | CGGTTCTGGTCRATCA | |

| | | | | |
|---|---|---|---|---|
| Comammox | comaA* | comaA-244F | TAYAAYTGGGTSAAYTA | Pjevac et al. 2017 |
| | | comaA-659R | ARATCATSGTGCTRTG | |
| | comaB | comaB-244F | TAYTTCTGGACRTTYTA | |
| | | comaB-659R | ARATCCARACDGTGTG | |
| Anammox | hzocl | hzocl1F1 | TGYAAGACYTGYCAYTGG | Li et al. 2010 |
| | | hzocl1R2 | ACTCCAGATRTGCTGACC | |
| | hzsA* | hzsA_1597F | WTYGGKTATCARTATGTAG | Harhangi et al. 2012; Oshiki et al. 2018 |
| | | hzsA1857R | AAABGGYGAATCATARTGGC | |
| DNRA | nrfA* | nrfAF2aw | CARTGYCAYGTBGARTA | Welsh et al. 2014 |
| | | nrfAR1 | TWNGGCATRTGRCARTC | |
| Denitrification | narG_W9F* | W9F | MGNGGNTGYCCNMGNGGNGC | Gregory et al. 2000 |
| | | T38R | ACRTCNGTYTGYTCNCCCCA | |
| | narG_1960f* | narG1960f | AYGTSGGSCARGARAA | Philippot et al. 2002 |
| | | narG2650r | TYTCRTACCABGTBGC | |
| | napA_V66 | V66 | TAYTTYYTNHSNAARATHATGTAYGG | Flanagan et al. 1999 |
| | | V67 | DATNGGRTGCATYTCNGCCATRTT | |
| | nirSC1F | nirSC1F | ATCGTCAACGTCAARGARACVGG | Wei et al. 2015a |
| | | nirSC1R | TTCGGGTGCGTCTTSAGAASAG | |
| | nirSC2F | nirSC2F | TGGAGAACGCCGGNCARGTNTGG | |
| | | nirSC2R | GATGATGTCCACGGCNACRTANGG | |
| | nirSC3F* | nirSC3F | TTCGCCCTGAARGAYGGNGG | |
| | | nirSC3R | AGGTGCCCACGAANARNCCNCC | |
| | nirK_FlaCu | FlaCu | ATCATGGTSCTGCCGCG | Throbäck et al. 2004 |
| | | R3Cu | GCCTCGATCAGRTTGTGGTT | |
| | nirK876 | nirK876 | ATYGGCGGVAYGGCGA | Henry et al. 2004 |
| | | nirK1040 | GCCTCGATCAGRTTRTGGTT | |
| | nirKC2F* | nirKC2F | TGCACATCGCCAACggnatgtwygg | Wei et al. 2015a |
| | | nirKC2R | GGCGCGGAAGATGshrtgrtcnac | |

| | | | | |
|---|---|---|---|---|
| | norB2 | norB2 | GACAARHWVTAYTGGTGGT | Casciotti and Ward 2005 |
| | | norB6 | TGCAKSARRCCCCABACBCC | |
| | cnorB-2F | cnorB-2F | GACAAGNNNTACTGGTGGT | Braker and Tiedje 2003 |
| | | cnorB-6R | GAANCCCCANACNCCNGC | |
| | qnorB2F-5R | qnorB2F | GGNCAYCARGGNTAYGA | |
| | | qnorB5R | ACCCANAGRTGNACNACCCACCA | |
| | qnorB2F-7R* | qnorB2F | GGNCAYCARGGNTAYGA | |
| | | qnorB7R | GGNGGRTTDATCADGAANCC | |
| | nosZ1F | nosZ1F | WCSYTGTTCMTCGACAGCCAG | Henry et al. 2006 |
| | | nosZ1R | ATGTCGATCARCTGVKCRTTYTC | |
| | nosZ-F-1181 | nosZ-F-1181 | CGCTGTTCITCGACAGYCAG | Rich et al. 2003 |
| | | nosZ-R-1880 | ATGTGCAKIGCRTGGCAGAA | |
| | nosZ-II-F* | nosZ-II-F | CTIGGICCIYTKCAYAC | Jones et al. 2013 |
| | | nosZ-II-R | GCIGARCARAAITCBGTRC | |
| | nosZ912F | NosZ912F | CGTCCCCGGCCTCGTGTA | Sanford et al. 2012 |
| | | NosZ1853R | GAGCAGAAGTTCGTGCAGTAGTAGGG | |
| Fungal denitrification | nirKfF | nirKfF | TACGGGCTCATGTAYGTNSARCC | Wei et al. 2015b |
| | | nirKfR | AGGAATCCCACASCNCCYTTNTC | |
| N fixation | nifHF* | nifHF | AAAGGYGGWATCGGYAARTCCACCAC | Rösch et al. 2012 |
| | | nifHR | TTGTTSGCSGCRTACATSGCCATCAT | |

* These assays did not produce amplifications on more than half of the samples or did not have a standard curve with at least three points and an $r^2$ value >0.95; therefore, were not used for the downstream analyses.

**Table S3.** List of the NiCE chip assays used for downstream analyses and the associated $r^2$ values of the standard curves

| Target | Assay ID | Standard Curve $r^2$ value | |
|---|---|---|---|
| | | Chip 1 | Chip 2 |
| Total Archaea | Arch_16S | 0.9989 | 0.9935 |
| AOB (Gammaproteobacteria) | Gamo_F1R1 | 0.9948 | 0.9988 |
| | Gamo_F1R2 | 0.9956 | 0.9976 |
| | Gamo_F2R1 | 1 | 0.9966 |
| AOB (Betaproteobacteria) | Bac_amoA | 0.9992 | 0.9965 |
| AOB hao | hao | 0.9991 | 0.9987 |
| AOA | Arch_amoAF | 0.9985 | 0.9982 |
| | Arch_amoAFA | 0.9995 | 0.9993 |
| NOB (Nitrobacter) | nxrBF | 0.9935 | 0.9984 |
| NOB (Nitrospira) | nxrB169f | 0.9983 | 0.9997 |
| Comammox | comaB | 0.9991 | 0.9985 |
| Anammox | hzocl | 0.996 | 0.9866 |
| Denitrification | napA_V66 | 0.9861 | 0.9935 |
| | nirSC1F | 0.9998 | 0.9982 |
| | nirSC2F | 0.9977 | 0.9996 |
| | nirK_FlaCu | 0.9999 | 0.9917 |
| | nirK876 | 1 | 0.9979 |
| | norB2 | 0.9976 | 0.9971 |
| | cnorB-2F | 0.9594 | 0.9885 |
| | qnorB2F-5R | 0.9966 | 0.9993 |
| | nosZ1F | 0.9998 | 0.9988 |
| | nosZ-F-1181 | 0.9841 | 0.996 |
| | nosZ912F | 0.9901 | 0.9991 |
| Fungal denitrification | nirKfF | 0.9929 | 0.9989 |
